# Predicting prime editing efficiency across diverse edit types and chromatin contexts with machine learning

**DOI:** 10.1101/2023.10.09.561414

**Authors:** Nicolas Mathis, Ahmed Allam, András Tálas, Elena Benvenuto, Ruben Schep, Tanav Damodharan, Zsolt Balázs, Sharan Janjuha, Lukas Schmidheini, Desirée Böck, Bas van Steensel, Michael Krauthammer, Gerald Schwank

## Abstract

Prime editing is a powerful genome editing technology, but its efficiency varies depending on the pegRNA design and target locus. Existing computational models for predicting prime editing rates are limited by their focus on specific edit types and by omitting the local chromatin environment. In our study, we developed machine learning models that predict prime editing efficiencies across a wide range of edit types up to 15 bp (’PRIDICT2.0’) and in different chromatin contexts (’ePRIDICT’). Both models can be accessed at www.pridict.it.

## Main

The efficiency of prime editing can vary considerably across different target sites and is heavily influenced by the design of the prime editing guide RNA (pegRNA). Our team^1^ and other researchers^2–4^ have previously developed machine learning models trained on extensive prime editing datasets to predict pegRNA efficiencies. A shared limitation of these models is their specialization in predicting certain edit types: DeepPrime^4^ is limited to 1-3 bp edits, MinsePIE^3^ is confined to insertions, and PRIDICT^1^ primarily focuses on 1 bp replacements as well as short insertions and deletions. Furthermore, each of these models does not account for the potential influence of the local chromatin state on editing rates. In this study, we addressed these shortcomings by developing two complementary computational models. ‘PRIDICT2.0’ predicts prime editing efficiency across a wide spectrum of edit types, and ‘ePRIDICT’ (epigenetic-based PRIme editing efficiency preDICTion) assesses the influence of locus-specific chromatin features on editing rates.

To this end, we constructed a highly diverse target-matched pegRNA library (Library-Diverse), which includes 1-5 bp replacements, 1-15 bp insertions and deletions, and pairs of simultaneously encoded single base replacements at variable distances (**ExtFig1a,b**). Given that the mismatch-repair (MMR) pathway influences prime editing rates^5,6^, we conducted our screens in both MMR-deficient (HEK293T)^7^ and MMR-proficient (K562)^8^ cells (**Fig1a**, **ExtFig1c**). Cells with the stably integrated Library-Diverse were transfected with plasmids expressing the prime editor, followed by deep amplicon sequencing to analyze editing rates with different pegRNAs. Demonstrating the robustness of the generated datasets, we observed high correlations in editing efficiencies between replicates (Spearman (R)=0.97/Pearson (r)=0.99 for HEK293T and R=0.84/r=0.96 for K562; **ExtFig1d,e**). Next, we assessed differences in the efficiency of installing diverse edit types in HEK293T and K562 cells. In line with the hypothesis that MMR has a negative effect on short edits, we observed contrasting editing patterns between the two cellular contexts. In HEK293T, the efficiency of installing insertions gradually declined with increasing lengths (**Fig1b**), whereas in K562 cells, editing rates were most efficient for insertions with a length of 4-5bp (**Fig1c**). Likewise, the length of replacements had minimal impact on the editing efficiency in HEK293T cells (**Fig1d**), while in K562 cells, 3-5 bp long replacements were more efficiently installed than 1-2bp replacements (**Fig1e**). Interestingly, for installing deletions, we observed a similar pattern as for insertions in HEK293T cells, with an inverse correlation between the edit length and efficiency (**Fig1f**), whereas in K562 cells editing rates remained consistently low regardless of the length of the deletion (**Fig1g**). We then investigated whether incorporating additional 1bp replacements to the intended edit could reduce MMR recognition and thereby elevate editing rates. While in our initial analysis we did not observe a notable difference in editing rates between single- and double edits in both HEK293T and K562 cells (**Fig1h,i**), a position-specific analysis revealed that introducing a co-edit within the GG PAM sequence substantially improved the editing efficiency of 1bp replacements that are situated outside of the PAM (**Fig1j,k**). Notably, we also observed that particularly in K562 cells double edits that are located further apart frequently lead to intermediate editing, where only one of the two edits is installed (**Fig1h,i**).

**Fig. 1.**
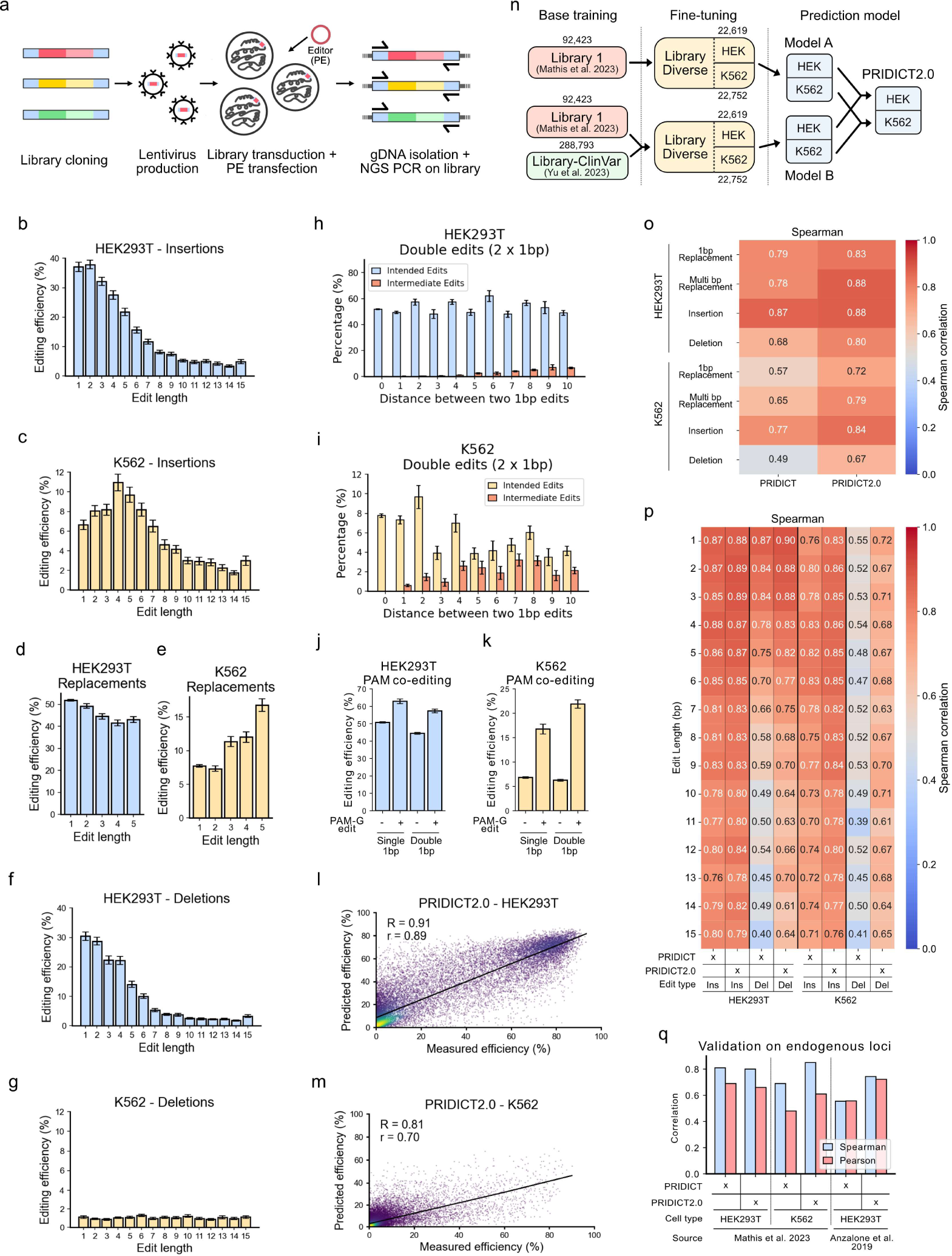
Characterization and prediction of pegRNA efficiencies based on sequence context. (**a**) Schematic overview of the screen with the target-matched pegRNA library ‘Library Diverse’. (**b**-**g**) Editing efficiency for (**b,c**) insertions in HEK293T or K562, for (**d,e**) 1-5bp replacements in HEK293T or K562, and (**f,g**) 1-15 bp deletions in HEK293T or K562. (**h,i**) Editing efficiencies in HEK293T (**h**) or K562 (**i**) cells for double edits where 2 separated 1 bp replacements were installed. Intended editing means that both replacements were installed, whereas intermediate editing means that only 1 of the 2 replacements was installed. Distance of 0 corresponds to single 1 bp edits. (**j,k**) Editing efficiency of single and double 1 bp replacements with or without editing within the GG PAM sequence in HEK293T (**j**) and K562 (**k**) cells. (**d**,**e,h-k**) Bars include only pegRNAs with 7, 10, or 15 bp RTT overhang to ensure similar RTT overhang distributions between conditions. (**b-k**) Bars show mean with error bar indicating mean +/− s.e.m. (**l**,**m**) Performance of PRIDICT2.0 on Library-Diverse (5-fold cross-validation) for (**l**) HEK293T (n = 22,619) and (**m**) K562 (n = 22,752) cells. Color gradient from dark purple to yellow indicates increasing point density, per Gaussian KDE. (**n**) Schematic illustration of PRIDICT2.0, which is an ensemble model that is based on the prediction average of two models: (Model A), base trained on Library 1^1^ and fine-tuned on Library-Diverse (HEK293T and K562), and (Model B), base trained on Library 1^1^ and Library-ClinVar^4^ and again fine-tuned on Library-Diverse. The number of pegRNAs in each dataset is indicated above or below the datasets. (**o**,**p**) Performance of PRIDICT vs. PRIDICT2.0 on different edit types (**o**) and insertions and deletions with different lengths (**p**). (**q**) Validation of PRIDICT2.0 on datasets where endogenous loci were targeted.

Next, we trained different machine learning models on the prime editing data generated with ‘Library-Diverse’ in HEK293T and K562 cells. Benchmarking their performance, we found that the attention-based bidirectional recurrent neural network (AttnBiRNN) model surpassed tree-based and linear regression models (**ExtFig2a-d**). To increase the robustness of the model to other experimental settings, we next augmented the training data with additional datasets obtained from other target-matched pegRNA library screens (’Library 1’^1^, ‘Library-ClinVar’^4^). The applied AttnBiRNN model extends on the architecture of our previous PRIDICT^1^ model and uses a simultaneous training strategy to predict prime editing efficiencies in HEK293T and K562 cells after fine-tuning on the ‘Library-Diverse’ datasets (**Fig1n**). The final model, termed PRIDICT2.0, was in total trained on over 400,000 pegRNAs and achieved a correlation of R=0.91/r=0.89 in HEK293T cells (**Fig1l**) and R=0.81/r=0.7 in K562 cells (**Fig1m**). PRIDICT2.0 thereby surpassed the performance of the original PRIDICT^1^ model, which achieved an R=0.82/r=0.78 in HEK293T cells and an R=0.63/r=0.40 in K562 cells on the same ‘Library-Diverse’ test dataset. A more detailed analysis revealed higher correlations for PRIDICT2.0 across all editing types and in both HEK293T and K562 cells, with the performance increase being more pronounced in K562 cells (**Fig1o,p**, **ExtFig2e,f**).

For further validation, we applied PRIDICT2.0 on a previously established prime editing dataset, where edits were introduced at different endogenous loci in HEK293T and K562 cells^1^. As the majority of edits in this dataset were 1bp replacements, the performance of PRIDICT2.0 in HEK293T cells was similar to PRIDICT, with R/r values of 0.8/0.66 vs. R=0.81/r=0.69^1^ (**Fig1q**). However, in K562 cells, PRIDICT2.0 outperformed PRIDICT, achieving an R=0.85/r=0.61 (**Fig1q**) compared to R=0.69/r=0.48^1^. Supporting our assumption that PRIDICT2.0 performs robustly under a wide range of experimental conditions, it also surpassed PRIDICT on a prime editing dataset generated in HEK293T cells by Anzalone et al.^9^ (PRIDICT2.0: R=0.74/r=0.72 vs. PRIDICT R=0.55/r=0.56^1^; **Fig1q**). In summary, PRIDICT2.0 is a versatile and robust deep learning model capable of predicting the editing efficiency of different edit types up to 15bp in mismatch-repair deficient and proficient contexts.

Previous research using SpCas9 has shown that genome editing rates are influenced not only by the sequence of the target locus but also by the chromatin environment^10^. To systematically explore a potential influence of chromatin on prime editing, we utilized the TRIP (Thousands of Reporters Integrated in Parallel) technology^11^ (**Fig2a, b**). A 640 bp TRIP reporter construct, which was shown to adapt the local chromatin environment^10^, was inserted into K562 cells using Piggy-Bac transposition and mapped by tagmentation PCR followed by NGS. Integration sites were widely distributed across all chromosomes (**Fig2c**) and most frequently mapped to genic regions (**Fig2d**). In the next step, we retrieved 455 publicly available ENCODE^12^ datasets for K562 cells, which include information on chromatin modification/accessibility and transcription factor binding (**Fig2e**). For each location, we averaged signals across different-sized windows from 100 to 5000 bp up and downstream of each location (**Fig2f**). We then transfected the cell pool with plasmids expressing the prime editor together with a pegRNA targeting the integrated construct (1182 mapped locations). To assess potential differences in chromatin effects to other commonly used genome editing techniques, we separately treated cells with an adenine base editor (ABE8e; 1169 mapped locations), cytosine base editor (BE4max; 1194 mapped locations), and a conventional Cas9 nuclease (SpCas9; 1196 mapped locations). By first adjusting the experimental conditions, we ensured that the editing rates followed a bell-curved distribution, with mean values ranging from 40% to 70% (**ExtFig3g-j**). Confirming the robustness of the experimental setup, we observed a strong correlation in editing efficiencies between individual screening replicates, with a Pearson (r) correlation of >0.93 for PE, >0.85 for ABE8e, >0.95 for BE4max and >0.87 for Cas9 (**ExtFig3a-d**). Notably, comparing different genome editors also revealed a strong correlation of PE with ABE8e (R=0.69/r=0.76) and BE4max (R=0.72/r=0.78), but only a moderate correlation with Cas9 (R=0.4, r=0.44) (**ExtFig3e-f**).

**Fig. 2.**
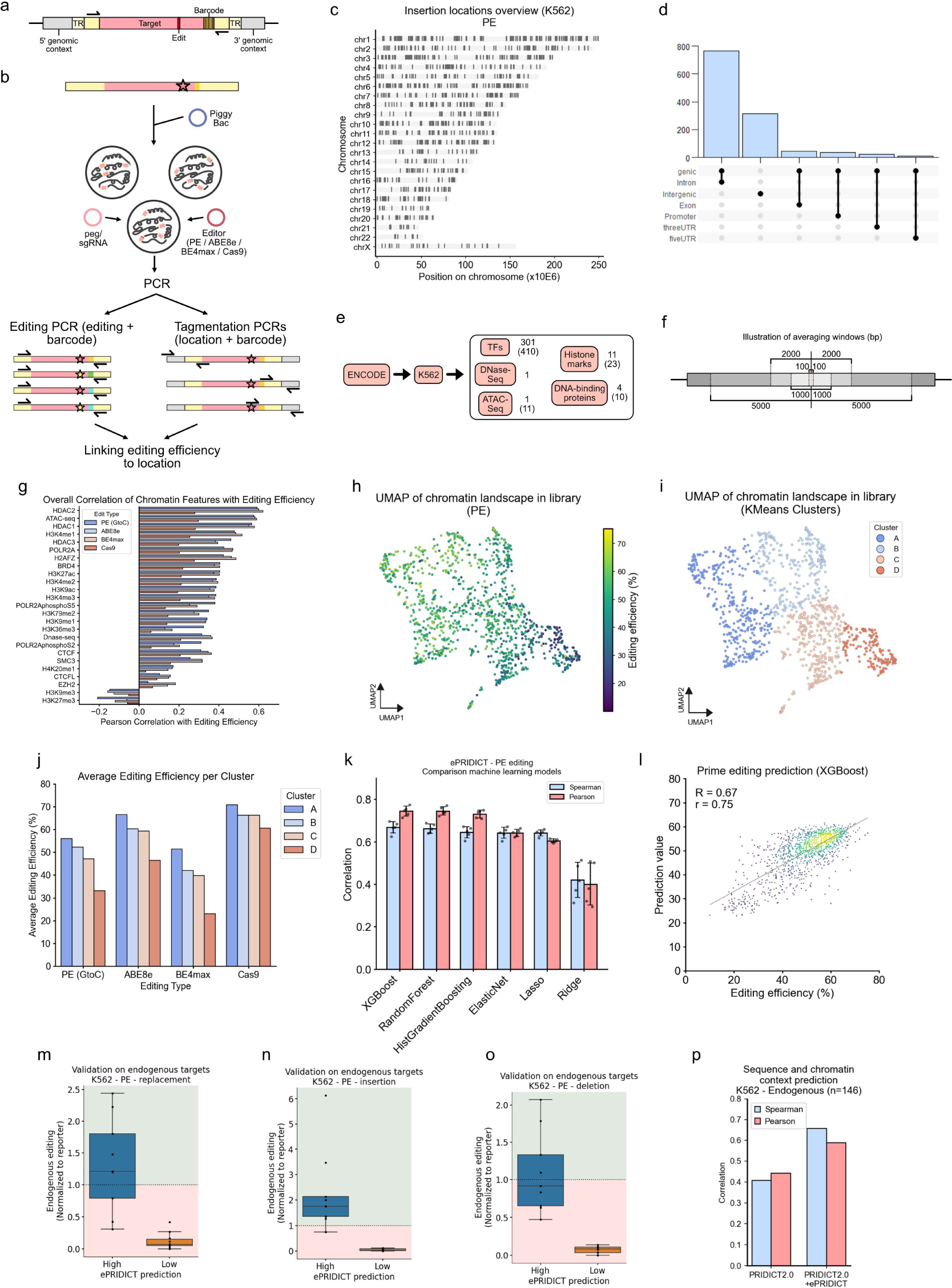
Characterization and prediction of prime editing efficiency in different chromatin contexts. (**a**) Schematic illustration of the TRIP library integrated by PiggyBac. TR: Terminal Repeats. (**b**) Schematic illustration of the TRIP screen in K562 cells. (**c**) Overview of TRIP reporter insertion locations with mapped prime editing efficiencies in the K562 genome. n = 1182. (**d**) Context of the TRIP reporter integration sites. (**e**) Schematic illustration of all ENCODE datasets of K562 used in this study. TF: Transcription Factors. The number of different features is indicated, with the total number of datasets (accounting for multiple ENCODE contributions per feature) given in brackets. Total number of datasets: 455. (**f**) Illustration of averaging windows (100, 1000, 2000, and 5000 up and downstream) around mapped integrations over which datasets are averaged for further analysis. (**g**) Overall Pearson correlation of a selection of chromatin characteristics (25/455) to editing efficiency (PE/prime editing, ABE8e, BE4max, and Cas9) across the TRIP library. The averaging window with the highest absolute correlation value with PE editing is shown for each feature. (**h**) UMAP projection based on all 455 ENCODE datasets and averaging windows of the TRIP library. Prime editing efficiency is shown via color scale. n = 1165. (**i**) KMeans clustering on UMAP projection to cluster integrations into 4 groups (A to D). n = 1165. (**j**) Average editing efficiency of integrations in each KMeans cluster for PE, ABE8e, BE4max, and Cas9. n per cluster: 380 (A), 267 (B), 349 (C), 169 (D). (**k**) Comparison of machine learning model performances on editing efficiency prediction with ePRIDICT on the TRIP library in K562 cells. Bars show the mean of fivefold cross-validation, and each of the five cross-validations is visualized as individual data points (n=5). Error bar indicates the mean +/− s.d. (**l**) Visualization of ePRIDICT XGBoost model predictions (n = 1182). Predictions from 5 cross-validations were combined for visualization. Color gradient from dark purple to yellow indicates increasing point density, per Gaussian KDE. (**m**-**o**) Validation of prime editing efficiency on endogenous loci with a high (>50) or low (<35) ePRIDICT score normalized to editing on the reporter sequence for (**m**) 1bp replacements (n-high: 8, n-low: 10), (**n**) 4bp insertions (n-high: 9, n-low: 9) and (**o**) 4bp deletions (n-high: 9, n-low: 10). (**p**) Performance of PRIDICT2.0 alone or in combination with ePRIDICT (average of prediction values from both models) on 41 endogenous loci with highly variable chromatin characteristics, targeted with 146 pegRNAs in K562 cells.

When we next analyzed the correlation between frequently studied chromatin features and editing efficiency, we observed that features correlating with open chromatin or active genes (such as ATAC-seq, HDAC1/2/3, or H3K4me1/2/3)^13–15^ positively correlated with editing rates (**Fig2g**). In contrast, repressive marks (H3K9me3, H3K27me3)^16,17^ were negatively correlated. For a more detailed analysis of different chromatin contexts, we performed UMAP projection using the extracted chromatin features of mapped genomic locations. When overlaying this projection with editing efficiency, we observed a low to high efficiency gradient, with locations resembling each other’s chromatin landscape (closer in the UMAP) having similar efficiencies (**Fig2h**, **ExtFig4a-c**). Further analysis of locus characteristics of the UMAP projection by grouping locations into 4 clusters (K-means; **Fig2i**) showed a similar pattern for different editors (PE, ABE8e, BE4max, and Cas9), with cluster A having the highest efficiency and cluster D the lowest (**Fig2j**). Cluster A was enriched in the active marks H3K4me3, H3K4me2, and H3K27ac, while showing a reduction in the repressive mark H3K27me3. This profile resembled a promoter-like environment^15^ (**ExtFig4d**). Cluster B had elevated levels of H3K36me3 and POLR2A and was reduced in the active mark H3K4me3, suggesting characteristics of transcriptional elongation within gene bodies^18^ (**ExtFig4e**). Cluster C showed increased levels of repressive H3K27me3 and CTCF and was reduced in active marks H3K4me3 and H3K27ac, indicative of repressed or insulated chromatin^16^ (**ExtFig4f**). Cluster D, with the lowest editing efficiency, had a slight elevation in the repressive mark H3K9me3 and was notably depleted in both active and elongation-associated marks, pointing to a more heterochromatic or repressed state^17^ (**ExtFig4g**).

In a subsequent step, we trained an array of linear and tree-based machine learning models to predict the influence of chromatin characteristics on prime editing efficiency (**Fig2k**). The model showing the highest correlations between predicted and experimentally observed editing rates is based on the XGBoost framework (**Fig2l**, R = 0.67, r=0.75) and was termed ‘ePRIDICT’ for epigenetic-based PRIme editing efficiency pre-DICTion. To next optimize the model for enhanced computational efficiency, we developed ePRIDICT-light, which is again based on the XGBoost framework but trained exclusively on 6 ENCODE datasets that showed high feature importance in ePRIDICT (HDAC2, H3K4me1, H3K4me2, H3K9me3, H3K27me3, and DNase-seq). Despite requiring fewer computational resources, ePRIDICT-light performed with similar accuracy to ePRIDICT, achieving R = 0.65 and r = 0.73 (**ExtFig4h**). Of note, we also developed XGBoost models for base editing and Cas9-mediated genome editing. While the base editor models showed similar performance as ePRIDICT (R = 0.69/r = 0.75 for ABE8e and R = 0.7/r = 0.75 for BE4max), the performance of the Cas9 model was substantially lower (R = 0.33/r = 0.35) (**ExtFig4i**).

For further validation of ePRIDICT, we also applied the model on a recently generated independent prime editing dataset^19^. Similar to our study, Li et al.^19^ performed prime editing on a single target site that was integrated into different genomic loci in K562 cells. Despite introducing another edit type (insertion instead of transversion) and targeting a different sequence, ePRIDICT reached a correlation of R=0.53, r=0.51 (**ExtFig4j**). Next, we validated ePRIDICT on endogenous loci. We selected 19 genomic sites with varying ePRIDICT values and integrated cassettes containing these sequences into K562 cells via PiggyBac transposition (MOI > 1). Treatment of these cell lines with the prime editor and locus-specific pegRNAs allowed us to directly compare prime editing rates on endogenous loci vs. the identical sequences integrated randomly via Piggy-Bac transposition. For each locus, we tested three edit types (1bp replacement, 4bp insertion, and 4bp deletion). Importantly, we found that loci with high ePRIDICT scores (>50) showed higher editing rates at endogenous loci compared to reporter loci, whereas loci with low ePRIDICT scores (<35) showed lower editing at endogenous loci (**Fig2m-o**). Subsequently, we performed the identical experiment in HEK293T cells, which exhibit a similar chromatin landscape to K562 cells at the 19 targeted loci (**ExtFig5a**). Suggesting that ePRIDICT could also be applied to other cell types in case the chromatin landscape at the target region is comparable, we again observed higher relative editing rates at loci with high-vs. low ePRIDICT scores (**ExtFig5b-d**). Furthermore, a similar pattern in editing rates between endogenous loci and integrated cassettes was observed when the K562 and HEK293T cell lines were treated with base editors or Cas9 (**ExtFig5e-j**), indicating that ePRIDICT also provides insights about the targetability of endogenous loci with these genome editors.

Finally, we investigated whether ePRIDICT could elevate the performance of PRIDICT2.0 when loci with highly varying chromatin contexts are targeted. Therefore, we applied both models on a dataset where 41 of such loci were targeted by 146 pegRNAs in K562 cells (64 replacements, 42 insertions, 40 deletions). Importantly, the initially moderate performance of PRIDICT2.0 (R=0.41/r=0.44) was notably improved to R=0.66/r=0.59 when the model was combined with ePRIDICT (**Fig2p**, **ExtFig5k,l**). Likewise, in a dataset where 19 endogenous loci were targeted by 56 pegRNAs in HEK293T cells, the performance of PRIDICT2.0 increased when it was applied in conjunction with ePRIDICT (from R=0.44/r=0.46 to R=0.67/r=0.67; **ExtFig5m**).

In this study, we utilized target-matched library screening and TRIP to generate comprehensive prime editing datasets, enabling us to examine the influence of pegRNA design and local chromatin context on prime editing rates. Our target-matched library featured a diverse and equally distributed array of edit types, including single- and multi-base replacements, as well as insertions and deletions of varying sizes. This diversity facilitated the development of PRIDICT2.0, an exceptionally versatile model capable of accurately predicting edits up to 15 bp in length. Importantly, PRIDICT2.0 is also able to predict double edits, allowing researchers to assess if the co-introduction of a second mutation increases the editing rates of the intended edit. Analysis of our dataset revealed that this is often the case if the intended edit is located outside of the PAM and the GG PAM sequence is co-edited. Since PRIDICT2.0 was trained on prime editing data generated in HEK293T and K562 cells, the model allows the prediction of pegRNA efficiency in MMR proficient- or deficient contexts. As a general guideline, we recommend users to employ the ‘MMR-deficient’ HEK293T PRIDICT2.0 model for PE4^5^ approaches, in which the MMR pathway is inhibited during editing, and the ‘MMR-proficient’ K562 PRIDICT2.0 model for regular PE2 prime editing approaches in primary cells or cell lines with functional MMR.

Adapting TRIP for prime editing allowed us to systematically link chromatin characteristics with editing rates, leading to the generation of ePRIDICT, a computational model that provides a quantitative value for the editability of any specified locus in K562 cells. Analysis of our TRIP data, moreover, indicates that prime editing is more efficient in areas marked by chromatin features linked to promoter regions and actively transcribed genes. Conversely, editing is less effective in areas related to suppressed chromatin and heterochromatin. These insights provide an important reference for researchers aiming to gauge the potential of genetically modifying a genomic locus by prime editing.

In conclusion, we identified pegRNA and chromatin features that influence prime editing efficiency, and developed machine-learning models to guide researchers in designing their prime editing experiments. PRIDICT2.0 and ePRIDICT are both freely available via GitHub or at www.pridict.it.

**Extended Data Fig. 1.**
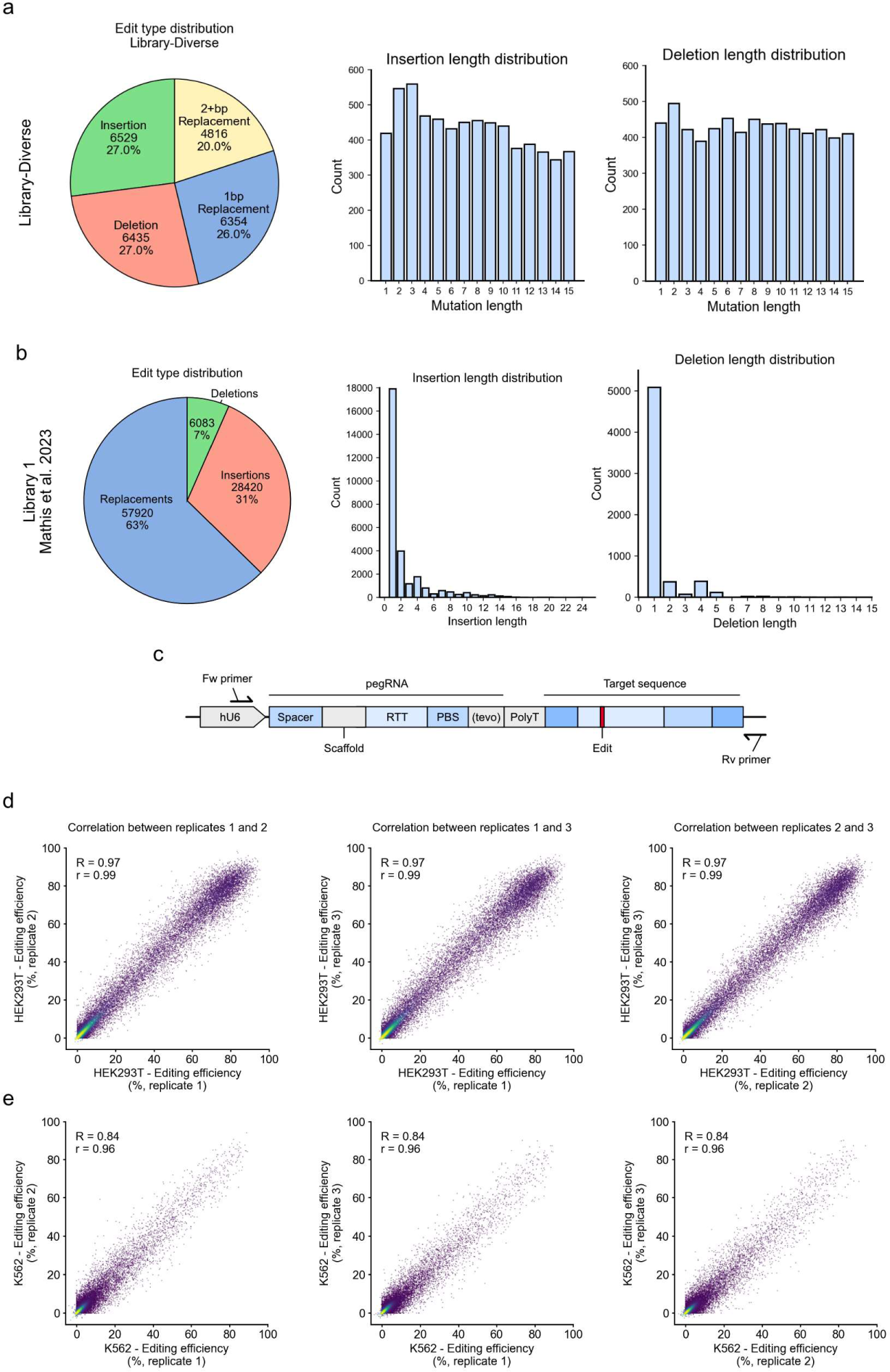
Library-Diverse characteristics. (**a**) Edit type distribution in ‘Library-Diverse’. (**b**) Edit type distribution in Library 1 from Mathis et al. 2023^1^, which has a focus on 1bp replacements and short insertions and deletions. (**c**) Self-targeting construct with the promoter (hU6) and different pegRNA domains (spacer, scaffold, reverse transcription template/RTT, primer binding sequence/PBS, tevopreQ1 motif, and poly T stop signal), target sequence, and primer location for NGS-PCR (forward (Fw) and reverse (Rv) primer). (**d, e**) Correlation of individual replicates of ‘Library-Diverse’ prime editing screens in (**d**) HEK293T (n = 22,619) and (**e**) K562 cells (n = 22,752). Color gradient from dark purple to yellow indicates increasing point density, per Gaussian KDE.

**Extended Data Fig. 2.**
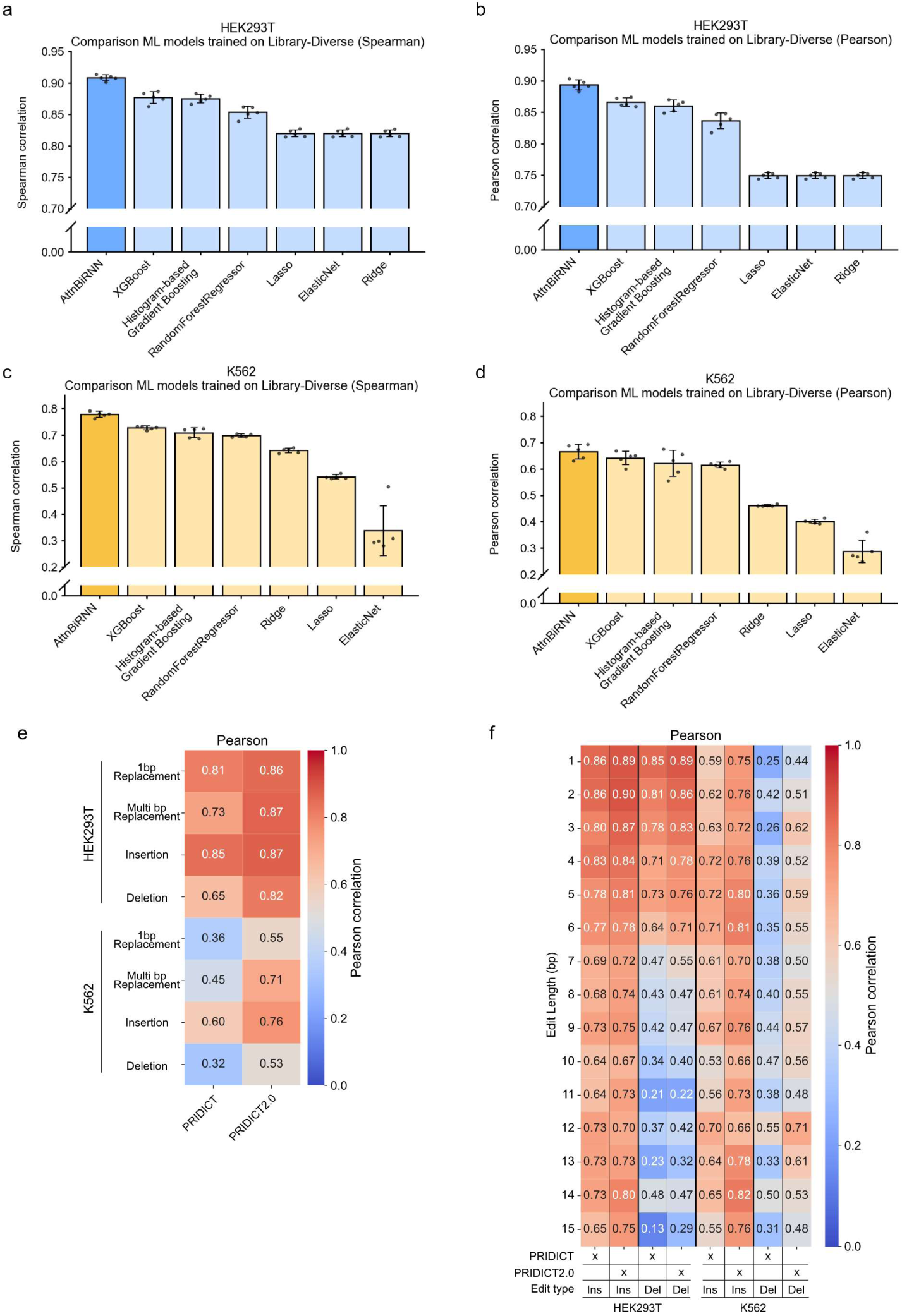
Machine learning metrics for training new models with ‘Library-Diverse’. (**a**, **b**) Comparison of machine learning model performances on editing efficiency prediction in HEK293T (Spearman (**a**), Pearson (**b**)) and K562 (Spearman (**c**), Pearson (**d**)). Bars show the mean of fivefold cross-validation, and each of the five cross-validations is visualized as individual data points (n=5). Error bar indicates the mean +/− s.d. (**e**) Prediction Pearson correlations of PRIDICT^1^ and PRIDICT2.0, tested on different edit types and cell types. (**f**) Prediction Pearson correlations of PRIDICT^1^ and PRIDICT2.0, tested on insertions and deletions of different lengths in HEK293T and K562 cells.

**Extended Data Fig. 3.**
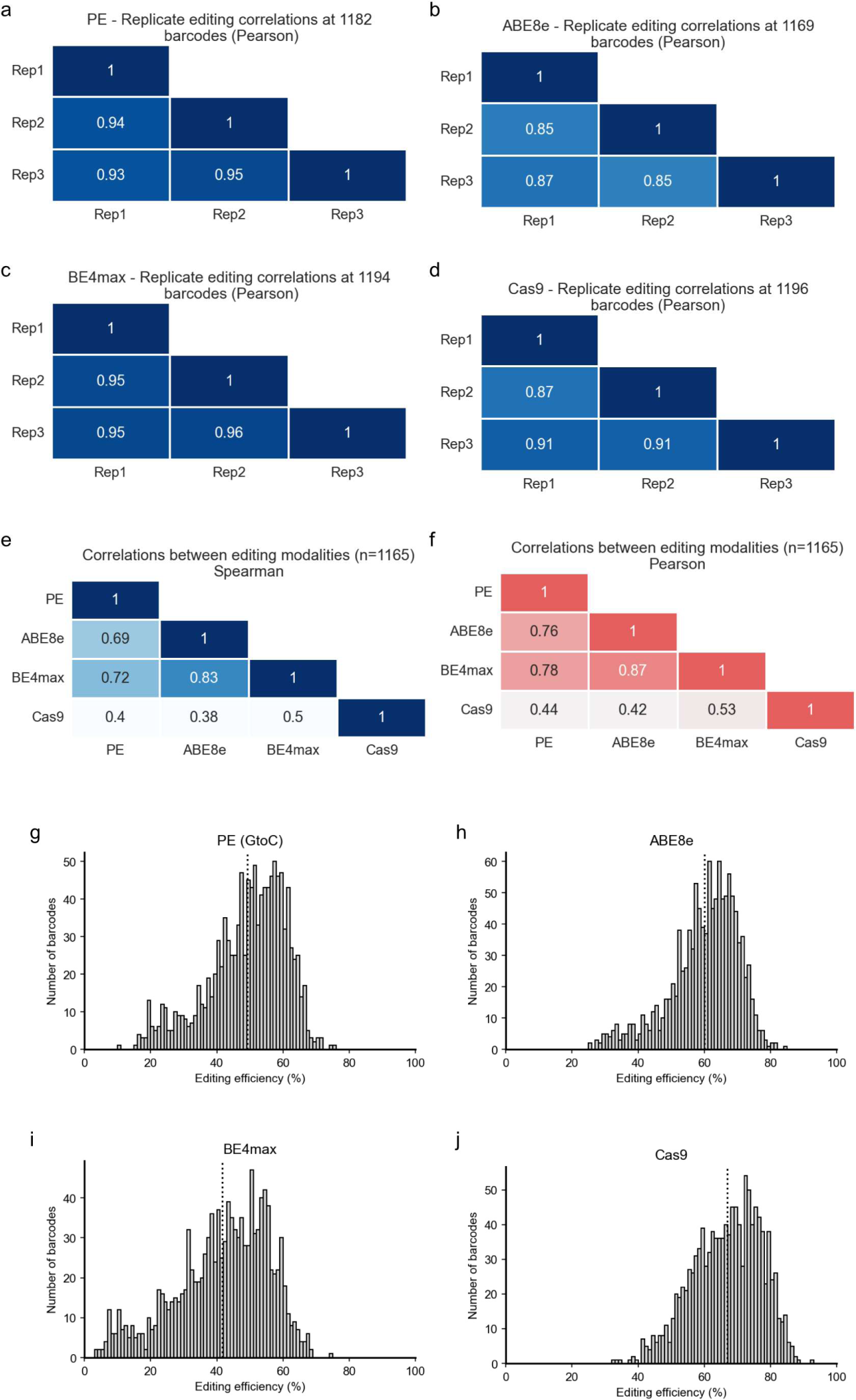
TRIP screen characteristics for different edit modalities. (**a-d**) Correlation (Pearson) of individual TRIP screening replicates for PE (n = 1182) (**a**), ABE8e (n = 1169) (**b**), BE4max (n = 1194) (**c**), and SpCas9 (n = 1196) (**d**) genome editing. (**e**,**f**) Correlation of replicate means between different edit modalities: Spearman (**e**), Pearson (**f**). Only barcode integrations available from all editors are used for analysis (n = 1165). (**g**-**j**) Distribution of editing efficiency across different TRIP reporter integrations for (**g**) PE, (**h**) ABE8e, (**i**) BE4max, and (**j**) SpCas9 genome editing. Dotted vertical line indicates mean editing efficiency.

**Extended Data Fig. 4.**
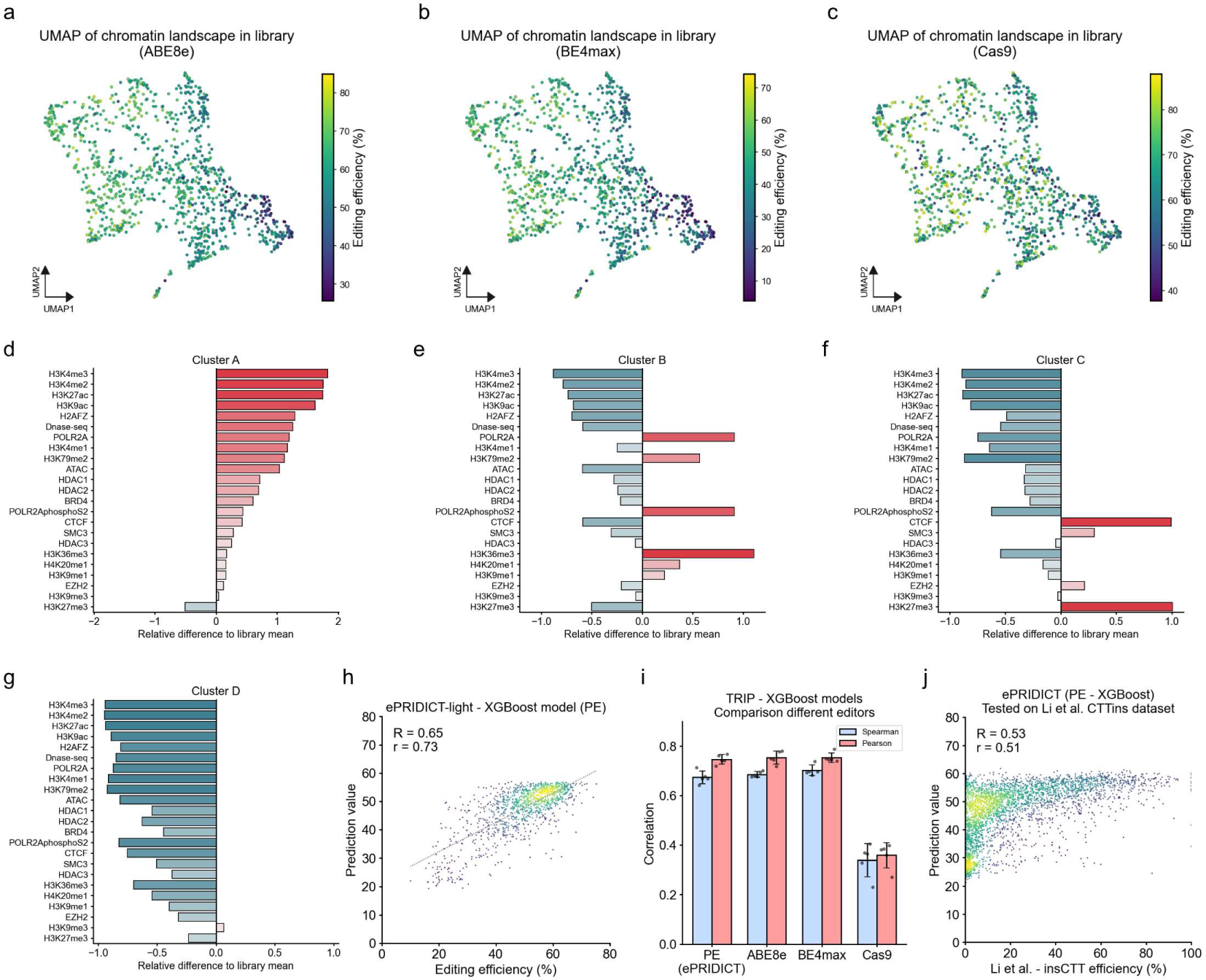
Additional analysis of TRIP screens and predictive modeling of editing rates. (**a**-**c**) UMAP projection based on chromatin characteristics of genomic locations in the TRIP library (n=1165; corresponding to integrations with mappings to all editors), with editing efficiency overlay of (**a**) ABE8e, (**b**) BE4max, and (**c**) Cas9. (**d**-**g**) Visualization of chromatin characteristics of clusters defined in **Fig2i**. For each target/dataset type, we selected the averaging window with the largest deviation from the library mean. The relative difference to the library mean, calculated as the absolute difference between the cluster average and the library mean divided by the library mean is shown. (**h**) Evaluation of the ePRIDICT-light XGBoost model trained on a subset of 6 features. Predictions from 5 different cross-validation runs were combined. (n=1182). (**i**) Spearman and Pearson correlation of XGBoost model prediction to editing efficiencies in TRIP library for prime editing, adenine base editing (ABE8e), cytosine base editing (BE4max), and SpCas9 genome editing. (**j**) Validation of ePRIDICT on an independent dataset from Li et al.^19^, where one sequence was integrated and edited (CTT insertion) at 4144 genomic locations. (**h**,**j**) Color gradient from dark purple to yellow indicates increasing point density, per Gaussian KDE.

**Extended Data Fig. 5.**
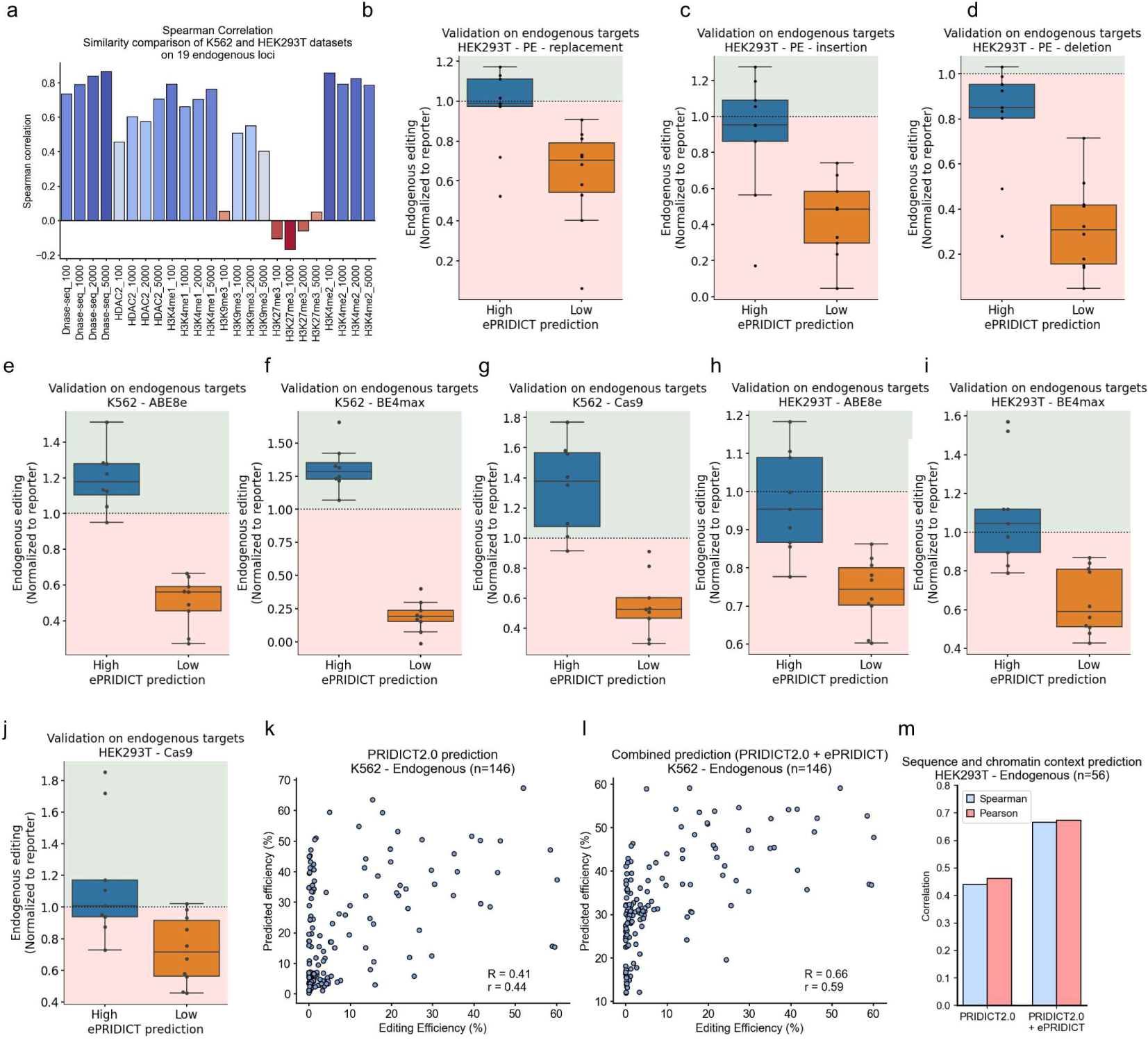
Validation of ePRIDICT at endogenous loci in K562 and HEK293T cells. (**a**) Spearman correlation analysis of ENCODE feature values for 19 selected endogenous loci, comparing datasets from K562 and HEK293T cells. (**b**-**d**) Validation of prime editing efficiency in HEK293T cells on endogenous loci with high (>50) or low (<35) ePRIDICT scores, normalized to editing on the reporter sequence. (**b**) 1bp replacements (n-high: 8, n-low: 10), (**c**) 4bp insertions (n-high: 9, n-low: 9), and (**d**) 4bp deletions (n-high: 9, n-low: 10). (**e-g**) Validation of genome editing efficiency on endogenous loci with high (>50, n = 8) and low (<35 n = 9) ePRIDICT values for (**e**) ABE8e in K562, (**f**) BE4max in K562, (**g**) Cas9 in K562, (**h**) ABE8e in HEK293T, (**i**) BE4max in HEK293T and (**j**) Cas9 in HEK293T, normalized to editing on the reporter sequence. (**k,l**) Performance of PRIDICT2.0 (**k**) or PRIDICT2.0 in combination with ePRIDICT (**l**) in predicting the editing efficiency of 146 pegRNAs targeting endogenous loci in K562. (**m**) Performance of PRIDICT2.0 alone or in combination with ePRIDICT (average of prediction values from both models) on 56 pegRNAs targeting endogenous loci in HEK293T, including highly and poorly accessible loci.

## Methods

### Clonings

The TRIP plasmid library (pPTK-BC-IPR) utilized in the chromatin-context investigation has been described in Schep et al. 2021^10^. Library-Diverse cloning was performed according to the protocol described by Mathis et al. 2023^1^. For the validation of ePRIDICT on endogenous targets, pegRNAs were designed as follows. We selected 20 genomic locations from the TRIP screen results, where 10 exhibited high editing efficiency for prime editing, ABE8e, and BE4max, and the remaining 10 showed low editing efficiency. For each location, a 200 bp window both upstream and downstream was scanned to identify pegRNAs capable of introducing a 1 bp replacement edit, a 4 bp insertion edit, and a 4 bp deletion edit. The pegRNAs with the highest scores based on PRIDICT^1^ were selected. Additionally, protospacers that contained at least one “A” for ABE8e and “C” for BE4max within their respective editing windows were identified. The highest-scoring spacer based on BEHive^20^ was selected to ensure the chosen gRNAs were highly predicted, minimizing the influence of sequence context on editing outcomes. Independently from these 20 previously described genomic locations, we designed 90 additional pegRNAs (21 deletions, 24 insertions, and 45 1bp replacements) targeting intronic and intergenic regions of 17 different genomic loci. All pegRNAs were cloned using established methods^1^, but with the pU6-tevopreq1-GG-acceptor^21^ vector. sgRNAs were incorporated into the lentiGuide-Puro^22^ (Addgene no. 52963) plasmid at BsmBI sites through one-pot cloning. Specifically, 5 μM of each spacer oligonucleotide were mixed in 1x Buffer 3.1, supplemented with 0.3 μl of BsmBI enzyme, 0.5 μl of T4 DNA ligase, 500 μM ATP, and 50 ng of the lentiGuide-Puro vector. This 20 μl reaction mixture was incubated at 37 °C for 1 hour before transformation. pegRNA and sgRNA plasmids were transformed into NEB Stable Competent E. coli. Resulting colonies were cultivated in LB (with 100 µg/ml carbenicillin) overnight. Plasmids were then extracted using the GeneJET Plasmid Mini-prep Kit (Thermo Fisher) and verified via Sanger sequencing.

### Oligo Library Design for Library-Diverse

Library-Diverse was built from two separate parts. First, 2,000 pathogenic ClinVar variants (1bp SNPs) were selected, and pegRNAs were designed to correct the sequence to wild-type (1). Next, a non-coding bystander mutation was determined for each variant, and two additional pegRNAs were designed, one with the correction edit and the bystander edit (2) and one with the bystander edit only (3). 942 pegRNAs were designed to have identical spacer sequences for all three pegRNAs (1-3) per target, while pegRNAs of the other targets had 2 or 3 different spacers (but identical correction edit). For this library, PBS length was kept constant at 13 bp and RTT overhang length at 10 bp.

For more flexibility in designing pegRNAs while preventing editing on endogenous sequences in the human genome, the second part of the library was designed based on the Emu (bird) genome. pegRNAs for 7,938 random deletions (1-15 bp), 7,941 random insertions (1-15 bp), 3,956 random multi-base changes (continuous 2-5 bp changes), and 3,968 random 1 bp changes were designed. PBS length was kept constant at 13 bp while RTT overhang varied (3, 7, 10, or 15 bp).

In total, the Library-Diverse was finally made up of 29,804 pegRNA-target combinations. The library was ordered from Twist Bioscience. Library oligos are listed in **Supplementary Table 1**.

### Cell culture

HEK293T (ATCC CRL-3216) were maintained in DMEM++ (DMEM plus GlutaMAX (Thermo Fisher Scientific), supplemented with 10% (vol/vol) fetal bovine serum (FBS, Sigma-Aldrich) and 1% penicillin-streptomycin (Thermo Fisher Scientific)) at 37 °C and 5% CO2. TrypLE Express (Thermo Fisher Scientific) was used for splitting HEK293T cells. K562 cells (ATCC CCL-243) were maintained in RPMI++ (RPMI 1640 Medium with GlutaMAX Supplement (Thermo Fisher Scientific) supplemented with 10% (vol/vol) fetal bovine serum (FBS, Sigma-Aldrich) and 1% penicillin-streptomycin (Thermo Fisher Scientific)) at 37 °C and 5% CO2. Cells were maintained at confluency below 90% and were tested negative for Mycoplasma contamination.

### PRIDICT2 library screen setup

#### HEK293T

HEK293T screening with Library-Diverse was performed as previously described (see “Library 1” in Mathis et al. ^1^). In short, we first produced lentivirus containing the library, then transduced HEK293T cells at an MOI < 0.3 and finally selected cells with 2.5 µg/µl puromycin for 7 days. After selection, cells were frozen and then thawed for each replicate independently. After thawing, cells were expanded and transfected (day 0) with PEI at a coverage of > 500x with pCMV-PE2-tagRFP-BleoR (Addgene no. 192508). One day later, cells were selected and maintained for 6 days with 750 ng/µl Zeocin (Invivogen) until harvest on day 7.

#### K562

K562 cell library with Library-Diverse was created as previously described^1^. In short, we first produced lentivirus containing the library, then transduced K562 cells at an MOI < 1, and finally selected cells with 2.5 µg/µl puromycin for 10 days. After selection, cells were frozen and then thawed for each replicate independently. pCMV-PE2-BleoR plasmid was electroporated (1450 V, 10 ms, 3 pulses) in the cell pool using a Neon Transfection System (100 µl kit). For each replicate, a total of 150 million K562 cells were transfected, with 3 million K562 cells and 10 µg of pCMV-PE2-tagRFP-BleoR (Addgene no. 192508) per individual electroporation (Day 0). Controls were electroporated without DNA. One day after electroporation (Day 1), the medium was replaced with RPMI++ with 200 ng/µl Zeocin. Selection was continued until collection day (Day 7).

### TRIP library screens

K562-TRIP cell library was generated with K562 cells (ATCC CCL-243), the piggyBac transposase (mPB-L3-ERT2.TatRRR-mCherry^11^), and a piggyBac plasmid library containing target sequence and barcode (pPTK-BC-IPR^10^). Electroporation was performed using a Neon Transfection System (100 µl kit) with 7.5 µg of editor plasmid and 2.5 µg guide plasmid per electroporation (Day 0; 3 million cells per electroporation; 1450 V, pulse width 10 ms, 3 pulses). pCMV-PE2-tagRFP-BleoR (Addgene no. 192508^1^) was used for prime editing, p2T-CMV-BE4max-BlastR (Addgene no. 152991^20^) was used for cytosine base editing, pCMV-ABE8e-SpG-P2A-GFP (in-house cloned from Addgene no. 138489^23^ and 140002^24^) was used for adenine base editing and pCMV-T7-SpCas9-P2A-EGFP (Addgene no. 139987^24^) for Cas9 editing. Guide sequences are listed in **Supplementary Table 1**. One day after electroporation (Day 1), the medium was replaced with RPMI++ with either 200 ng/µl Zeocin (PE) or 2.5 ng/µl puromycin (BE4max, ABE8e, Cas9). Cas9 cells were harvested the next day (Day 2), while PE, BE4max, and ABE8e cells were further selected until harvest on Day 7.

### Endogenous reporter validations

A reporter containing the genomic sequences of 10 highly scoring and 10 low scoring locations (see in Cloning methods part) from the TRIP libraries was designed and ordered as 3 separate gene blocks from IDT. Gibson assembly was performed to clone the reporter into a piggyBac-PuroR backbone. Next, HEK293T and K562 cells were transfected as follows. 130,000 HEK293T cells or 100,000 K562 cells were seeded into 48-well plate 6h before transfection and then transfected with 225 ng of reporter plasmid and 25 ng of piggyBac helper plasmid and 1 µl Lipofectamine 2000. Three days after transfection, the medium was replaced with DMEM++ and 2.5 ng/µl puromycin (HEK293T) or RPMI++ and 2.5 ng/µl puromycin (K562). Cells were selected for 10 more days to ensure the integration of the reporter construct.

HEK293T-reporter cells were then edited as follows. 130,000 cells were seeded per well in 48-well plate 6h before transfection (Day 0). Next, transfection was performed with 1 µl of Lipofectamine 2000 with:

PE: 750 ng of editor (pCMV-PE2-tagRFP-BleoR) and 250 ng of pegRNA
ABE8e, BE4max, Cas9: 500 ng of editor, 250 ng of sgRNA, 200 ng of pCMV-Tol2 (Addgene no. 31823) and 50 ng CMV-GFP. (ABE8e: p2T-CMV-ABE8e-SpCas9-BlastR, BE4max: p2T-CMV-BE4max-BlastR (Addgene no. 152991), Cas9: p2T-CMV-SpCas9-WT-BlastR)

Guide sequences are listed in **Supplementary Table 1**.

On the next day (Day 1), HEK293T cells were split 1:2 into new 48-well plates, and the medium was replaced with DMEM++ and 750 ng/µl Zeocin (PE) or 20 ng/µl Blasticidin (ABE8e, BE4max and Cas9). Cas9 transfected cells were harvested one day later (Day 2). PE, ABE8e, and BE4max transfected cells were maintained under selection until harvest on Day 7. K562-reporter cells were handled similarly but with the following differences. 100,000 cells were seeded per well, and selection was performed with RPMI++ and 200 ng/µl Zeocin (PE) or 20 ng/µl Blasticidin (ABE8e, BE4max and Cas9). Editing with the 90 pegRNAs targeting 17 loci (intronic and intergenic) was performed as described above but in wild-type K562 cells.

### Genomic DNA isolation and HTS

Genomic DNA from Library-Diverse was isolated by Blood & Cell Culture DNA Maxi Kit (Qiagen; 30 million cells per condition). Genomic DNA from the TRIP library editing experiments was isolated by DNeasy Blood & Tissue Kit (4 million cells per condition). DNA from endogenous editing experiments was isolated by direct lysis using direct lysis buffer: 10 µl of 4× lysis buffer (10 mM Tris-HCl pH 8, 2% Triton X-100, 1 mM EDTA, 1% freshly added proteinase K) was added to cells resuspended in 30 µl of PBS and incubated at 60 °C for 60 min and 95 °C for 10 min. Genomic DNA for the TRIP tagmentation experiment was isolated by direct lysis followed by Phenol-Chloroform DNA purification and ethanol precipitation.

Target sites of library-diverse were amplified by NEBNext Ultra II Q5 Master Mix (NEB, 26 cycles), and target sites from the TRIP library were amplified by GoTaq G2 Hot Start Green Master Mix (Promega, 26 cycles). Amplicons were purified via gel extraction using a NucleoSpin Gel and PCR Clean-up Mini kit (Macherey-Nagel). Illumina sequencing adapters were added with Phusion High-Fidelity DNA Polymerase (NEB; seven cycles) and purified by gel extraction. TRIP tagmentation library was prepared as described under the “Library analysis TRIP” methods section.

Endogenous target sites and reporter target sites for arrayed editing were amplified by GoTaq G2 Hot Start Green Master Mix (Promega, 30 cycles) and were purified with Sera-Mag Select (Merck). Illumina sequencing adapters were added with Q5 High-Fidelity DNA Polymerase (NEB; seven cycles) and purified by gel extraction.

Final pools were quantified on a Qubit v.3.0 (Invitrogen). The average amplicon size of the TRIP tagmentation library was quantified on a TapeStation 4200 (Agilent). Library-Diverse and TRIP editing and tagmentation libraries were sequenced paired-end on an Illumina NovaSeq 6000 using SP Reagent Kits (2×250 cycles for Library-Diverse, 2x 150 cycles for TRIP libraries). Arrayed targets (endogenous and reporter) were sequenced paired-end (2 × 150) on an Illumina MiSeq using MiSeq Reagent Micro Kit v2.

### Library analysis of Library-Diverse

Editing levels were determined by initial trimming of sequencing reads with Cutadapt v. 3.1^25^, followed by in-house scripts. Each sequencing read was assigned to the corresponding target sequence based on the spacer sequence, extension sequence, tevopreQ1 motif, and a barcode. Only reads with matches for all elements were used for the final analysis (filtering out ∼34% of total reads). To calculate the editing rate, we compared the read sequence (2 bp upstream of nick position until 5 bp downstream of edited flap end) with wild-type and edited sequence and assigned the labels ‘unedited’, ‘edited’, or ‘nonmatch’. Editing efficiency was calculated as previously described^1^ with the formula:

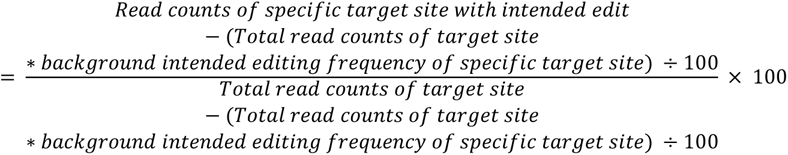

Background intended edit frequencies were determined by analyzing the control library pool, which was not transfected with the prime editor. Editing values were clamped to be within 0 and 100%. Target sequences were further filtered for having a minimum of 100 reads in every replicate. Target sequences where the corresponding control had >20% unintended editing rate or >5% intended editing were also discarded. This led to a total of 22,621 sequences for HEK293T and 22,752 sequences for K562 that were used for further analysis and machine learning. For each pegRNA, a selection of features was extracted to train statistical machine learning models and to complement deep learning models. These features included RTT-, PBS- and Correction-length, GC content, melting temperature, the maximum length of polyA/T/G/C sequence stretches, DeepSpCas9 score^26^, and minimum free energy (ViennaRNA Package v.2.0^27^). A full overview of features used in training statistical and deep learning models is listed in **Supplementary Table 2**.

### Library analysis TRIP

The tagmentation reaction was executed using the following protocol. To ensure high tagmentation coverage of the TRIP pool, 15 parallel reactions were conducted. Primers adapt_A & adapt_A_invT were annealed in a controlled ramp-down cycle ranging from 95 °C to 4 °C. The transposome complex was assembled by mixing 1 μl of 1:2 diluted adapters with 1.5 μl of Tn5 transposon in 18.7 μl of Tn5 dilution buffer (20 mM HEPES, 500 mM NaCl, 25% glycerol) and incubated for 1 hour at 37 °C. Tagmentation was performed by combining 100 ng of genomic DNA with 1 μl of the pre-assembled transposome and 2x TD buffer pH 7.6 in a 20 μl final volume. The reaction mixture was incubated at 55 °C for 10 minutes and subsequently quenched with 0.2% SDS. Three distinct libraries (“For,” “ForBC,” and “Rev”) were generated using linear PCR with For, ForBC, and Rev enrichment primers. The initial target enrichment involved 45 amplification cycles, using a mix that included tagmented DNA, primers, dNTPs, 5× Phusion HF Buffer, and Phusion HS Flex polymerase in a 20 µl final volume. For library preparation (PCR 1), 11 amplification cycles were conducted using Phusion HS Flex polymerase in a 25 µl final volume. N7xx adapters were introduced in PCR 2, consisting of 10 cycles of amplification using Phusion polymerase in a 22 µl final volume. Post-amplification, 5 µl of each reaction was assessed on a 1% agarose gel. The 15 parallel samples from the TRIP pool were combined. Libraries were bead-purified using CleanPCR beads at a 1:0.8 sample-to-bead ratio. Sequencing was done on a NovaSeq 6000 with 150 bp paired-end reads.

For sequencing analysis, the amplicons were trimmed by Cutadapt v. 3.1^25^ and aligned to the human genome with bowtie2 v. 2.5.1^28^ (hg38, mapq score >30). Only the ForBC mapping locations confirmed by both For and Rev amplicon mappings (with at least 10 mappings each), and those where locations of “For” and “Rev” locations were exactly 4bp from each other (TTAA integration motif of PiggyBac) were retained. Editing efficiencies of different editors and barcodes (= genomic insertions) were then analyzed by examining the target amplicon, which includes the target sequence and barcode. Following demultiplexing based on barcodes, editing efficiency for each barcode was computed using custom scripts. Control replicates were also evaluated, allowing for adjustments in the final editing rates to account for background editing/mutations. Genomic locations were then associated with editing efficiencies using the barcode sequence.

For correlating editing rates with chromatin characteristics, ChIP-seq, ATAC-seq, and DNase-seq, datasets for K562 were sourced from the ENCODE database^12^. All datasets are listed in **Supplementary Table 3**. Average values for different sequence lengths at genomic locations in our dataset were then computed. We selected windows of 100, 1000, 2000, and 5000 bp both upstream and downstream of each location, and averaged values across these regions (as shown in **Fig2f**). UMAP projection was employed to visually represent the chromatin landscape of our integrated reporter locations. Finally, K-means clustering was used to segment the locations into four distinct groups for a more in-depth analysis.

### Analysis of arrayed editing experiments

Arrayed experiments for validation on endogenous loci were analyzed using CRISPResso2 v. 2.2.12^29^ in batch mode. For prime editing samples, the original sequence (’amplicon_seq’), the expected sequence after editing (’expected_hdr_amplicon_seq’), and window of quantification (2 bp upstream of nick position until 5 bp downstream of edited flap end; ‘quantification_window_coordinates’) were used for batch analysis. For base editing and Cas9, the original sequence and guide sequence with default settings for base editing and a quantification window size of 3 for Cas9 were used. Editing efficiencies of 1-2 technical replicates (transfection in separate wells but on the same day) were averaged and used as one independent biological replicate for the following analysis. Control editing efficiencies were subtracted from each sample, and only samples with more than 500 reads were used in the analysis. Editing rates are listed in **Supplementary Table 1**.

### Machine learning of sequence- and chromatin-context-dependent editing efficiency

Machine learning models trained on Library-Diverse only (**ExtFig2a-d**) were developed based on 22,619 (HEK293T) and 22,752 (K562) pegRNAs. Training workflow for PRIDICT2.0 is depicted in a schematic illustration in **Fig1n**. Model A was built by base training with Library 1 (92,423 pegRNAs in HEK293T, from Mathis et al. 2023^1^) and then fine-tuning on Library-Diverse (with both HEK293T and K562 editing efficiencies). Model B was built by base training on Library 1 and Library-ClinVar (288,793 pegRNAs in HEK293T, from Yu et al. 2023^4^) and then fine-tuning on Library-Diverse. Finally, Model A and Model B were combined into an ensemble model, PRIDICT2.0, by combining prediction values of either HEK293T or K562 from both models in a 1:1 ratio. For all models, we followed a grouped fivefold cross-validation on Library-Diverse where pegRNAs for the same locus were kept in the same train- or test set. Each fold had 80% of the Library-Diverse pegRNAs for training and 20% for testing. A 10% grouped random split was taken from each fold’s training sequences to create a validation set, which was then used for optimizing the model’s hyperparameters. For the neural network model, we used a uniform random search strategy that randomly chose a set of hyperparameter configurations from the set of all possible configurations and trained^30^ corresponding models on a random fold. Subsequently, the best model hyperparameters were determined based on the performance of the models on the validation set of the respective fold. Finally, these hyperparameters were used for the final training and testing of each model on all five folds. For baseline models^31,32^ in **ExtFig2a-d** (XGBoost, Histogram-based Gradient Boosting, RandomForest Regressor, Lasso, ElasticNet, Ridge), we used a random search strategy over each model’s specific hyperparameter space where the best hyperparameters were determined using twofold cross-validation on the combined training and validation set of each fold. Subsequently, the model achieving the best performance was retrained and tested (using the test set) on each corresponding fold of the five folds.

For the chromatin context-based prediction of editing rates, we used the mapped editing efficiencies (1,182 for PE, 1,169 for ABE8e, 1,194 for BE4max, and 1,196 for Cas9) and ENCODE features from 455 datasets (ePRIDICT) or 6 datasets (ePRIDICT-light). We performed 5-fold cross-validation for PE prediction with linear regression-based models (Ridge, Lasso, ElasticNet) and tree-based models (HistGradientBoosting, RandomForest, XGBoost). Additionally, parameters for XGBoost ePRIDICT and ePRIDICT-light models were optimized via random search cross-validation (RandomSearchCV, Scikit-learn^31^). The same parameters were used for XGBoost models trained on ABE8e, BE4max, and Cas9 datasets. The performance of XGBoost models is shown by combining predictions from all individual folds and calculating correlations on the combined dataset. After benchmarking the model performances with cross-validation (**Fig2k,l,** and **ExtFig4h**), we retrained ePRIDICT and ePRIDICT-light on the full dataset (1,182 loci). For the predictions presented in **Fig2m-p** and **ExtFig5**, we trained an ePRIDICT model that excludes the loci featured in these figures from training to prevent training leakage.

### Analysis of third-party datasets

Editing rates for Library-ClinVar were extracted from the supplementary material of Yu et al.^4^. We incorporated all 288,793 pegRNAs, spanning both the Training and Test set, into the training process of the PRIDICT2.0 model. To validate ePRIDICT, we utilized the dataset from Li et al.^19^, which includes 4144 integration locations and the corresponding prime editing efficiencies. Feature values were extracted from all averaging windows (100, 1000, 2000, and 5000 bp up and downstream) across 455 ENCODE datasets, following the same methodology applied to our TRIP library.

### Statistics and reproducibility

Statistics were performed using Python v. 3.9.15 or 3.10.12 and SciPy v.1.10.1. Pearson (r) and Spearman (R) correlations were determined to evaluate the correlation of predicted and measured pegRNA editing rates. Editing experiments were performed at least in independent biological duplicates. Sample sizes, bars, box plots, and error bars are described in figure legends.

## Data availability

Measured editing rates used for analysis and creating figures in this study are provided in **Supplementary Table 1**. DNA-sequencing data will be added to the National Center for Biotechnology Information Sequence Read Archive upon publication. ENCODE datasets are listed in **Supplementary Table 3** and are available from encodeproject.org.

## Code availability

Scripts used in this study for data analysis or offline running of the prediction models (PRIDICT2.0, ePRIDICT) are provided on GitHub (github.com/uzh-dqbm-cmi/PRIDICT2, github.com/Schwank-Lab/epridict). Online implementation of both models can be accessed via www.pridict.it. Additional information on the PRIDICT2.0 algorithm can be found in **Supplementary Methods 1**.

## Acknowledgements

We thank the Functional Genomics Center Zurich for their support in next-generation sequencing; the Science IT team at the University of Zurich for the computational infrastructure used for data analysis; the ENCODE consortium for providing the datasets used for the analysis of TRIP libraries; C. Leemans for consulting during the TRIP library analysis; G. Affentranger for assistance in figure design; and the members of the Schwank laboratory for fruitful discussions. This work was supported by the Swiss National Science Foundation (SNSF) grant numbers 185293, 214936, and 201184, URPPs (University Research Priority Programs) ‘Human Reproduction Reloaded’ and ‘ITINERARE’, and the SERI financed ERC Consolidator Grant “GeneREPAIR”.

## Author contributions

N.M. designed the study, performed experiments, and analyzed data. A.A. designed and generated attention-based bidirectional RNNs (PRIDICT1.1 and PRIDICT1.2). Linear regression and tree-based machine learning models for Library-Diverse were built by A.A., and models for TRIP predictions were built by N.M.. E.B. and S.J. were involved in TRIP library screening. A.T. and T.D. performed editing experiments on endogenous loci. R.S. and B.v.S. provided the TRIP plasmid library and performed tagmentation experiments of the cell pool. Z.B. contributed to the integration analysis of the TRIP library. L.S. and D.B. helped in cloning experiments. N.M. and G.S. wrote the manuscript with input from A.A. and R.S.. M.K. and G.S. designed and supervised the research. All authors revised the manuscript.

