## Supplementary Methods 1 for "Predicting prime editing efficiency across diverse edit types and chromatin contexts with machine learning"

### Supplementary Methods 1a:

#### 1.1 PRIDICT Model Overview

Our model adapts the design of **PRIDICT** model [1] developed in PyTorch [2] comprising three encoders and one decoder neural network. Two separate encoder networks used bidirectional attention-based recurrent neural networks (RNN) to encode and learn representations from a pair of ***original*** and ***mutated*** sequences, and a third encoder network that used a feed-forward network to encode and learn a representation from a set of derived/computed ***sequence-level*** features (such as melting temperature, minimum free energy, etc.). These learned representations (i.e. fixed-length vectors) were mapped using a decoder network to learn a probability distribution on three outcomes associated with ***edited***, ***unedited***, and ***unintended edit*** proportions.

In this work, we introduce new model components and training strategy to the original **PRIDICT** model [1]. (1) Starting with the sequence pairs (i.e. ***original*** and ***mutated***), we align both sequences by introducing a new symbol '−' in case of insertions and deletions. The aligned sequence pair is used as the raw input to the model. (2) We introduce multi-task learning to model training where we jointly optimize a unified model on multiple datasets. The unified model has a set of common layers exploiting shared patterns across datasets, and separate layers that capture datasets-specific patterns. (3) Lastly, in conjunction with multi-task learning, we use fine-tuning by first adapting models with **PRIDICT** architecture pretrained on large datasets and then fine-tuned on our libraries.

The model's architecture is comprised of the following blocks: An (1) **Embedding block** that embeds the nucleotides (i.e. ACGT letters and the alignment symbol '−') and their corresponding position annotations (binary indicators such as being part of protospacer, PBS or RTT) from one-hot encoded representation to a dense vector representation.
An (2) **Encoder block** that contains a multilayer bidirectional gated recurrent units (GRU) [3, 4], that takes the embeded representations from the **Embedding block** and computes another representation based on left-to-right and right-to-left processing of the tokens in the sequence (i.e. uses context representation in both directions) when learning the new vector representation of the different tokens.
Lastly, an (3) **Attention** block comprising of (a) global context/query vector $\overline{q}_{global}$ (i.e. trainable parameters) that pools (using weighted averaging based on attention scores) the sequence of token-level representations to one representation (i.e. fixed-length vector), and (b) RTT query vector $\overline{q}_{RTT}$ to pool token-level representations belonging to the RTT (i.e. contiguous part of the sequence where mutation occurs) to a fixed-length vector representation.
For the ***sequence-level*** derived features, the model's encoder network is comprised of a mix of **multilayer feed-forward** networks, residual connections and layer-normalization operations to map the input features to a dense vector representation.
Lastly, the models' **decoder network** stacks the five computed representations from the three encoder networks and uses a series of **multilayer feed-forward** networks, residual connections and layer-normalization layers to compute a probability distribution vector on the three outcomes representing ***edited***, ***unedited*** and ***unintended edit*** proportions. A formal description of each component of the model is described in their respective sections below.

#### 1.2 Sequence encoder network

##### 1.2.1 Embedding block

Formally, given a design variant sequence (mutated or original sequence) $\underline{S} = [x_1, x_2, \cdots, x_T]$, a nucleotide at position $t$ is represented by 1-of-$K$ encoding where $K$ is the size of the set of all nucleotide letters in the data such that $x_t \in [0,1]^K$ and $T$ is the length of the sequence. An embedding matrix $W_e$ is used to map the input $x_t$ to a fixed-length vector representation (Eq. 1)

$$e_t = W_e x_t \tag{1}$$

where $W_e \in \mathbb{R}^{d_e \times K}$, $e_t \in \mathbb{R}^{d_e}$, and $d_e$ is the dimension of vector $e_t$.
Similarly, each position $p_t$ in the sequence $\underline{S}$ has multiple set of annotations indicating if it belongs to protospacer, PBS, and RTT (denoted by $p_t^{proto}$, $p_t^{PBS}$, and $p_t^{RTT}$ respectively). These annotations are represented by binary indicators where $p_t^{proto}$, $p_t^{PBS}$, and $p_t^{RTT} \in [0,1]$. An embedding matrix corresponding to each annotation

$W_{p^{proto}}$, $W_{p^{PBS}}$, $W_{p^{RTT}}$ is used to map the binary indicators to a fixed-length vector representation (Eq. 2)

$$\begin{aligned}
a_t^{proto} &= W_{p^{proto}} p_t^{proto} \\
a_t^{PBS} &= W_{p^{PBS}} p_t^{PBS} \\
a_t^{RTT} &= W_{p^{RTT}} p_t^{RTT}
\end{aligned} \tag{2}$$

where $W_{p^{proto}}$, $W_{p^{PBS}}$, $W_{p^{RTT}} \in \mathbb{R}^{d_a \times 2}$, $a_t^{proto}$, $a_t^{PBS}$, $a_t^{RTT} \in \mathbb{R}^{d_a}$ and $d_a$ is the annotation embedding dimension.

Computed embeddings at each position were concatenated $\bigoplus$ (Eq. 3) to get a unified representation for every token/element in the sequence $\underline{S}$ (i.e. compute a new sequence $\underline{U} = [u_1, u_2, \cdots, u_T]$ where $u_t \in \mathbb{R}^{d_u}$, $\forall t \in [1, \cdots, T]$ and $d_u = d_e + 3 \times d_a$ for the original sequence and $d_u = d_e + 2 \times d_a$ for the mutated sequence).

$$u_t = [e_t \bigoplus a_t^{proto} \bigoplus a_t^{PBS} \bigoplus a_t^{RTT}] \tag{3}$$

##### 1.2.2 Encoder block: bidirectional GRU

The computed embeddings of sequence $\underline{U} = [u_1, u_2, \cdots, u_T]$ from the embedding block is further processed using a recurrent neural network (RNN) encoder block. A vanilla RNN will compute a hidden vector at each position (i.e. state vector $h_t$ at position $t$), representing a history or context summary of the sequence using the input and hidden states vector form the previous steps. Equation 4 shows the computation of the hidden vector $h_t$ using the input $u_t$ and the previous hidden vector $h_{t-1}$ where $\phi$ is a non-linear transformation such as $ReLU(z) = max(0, z)$ or $tanh(z) = \frac{e^z - e^{-z}}{e^z + e^{-z}}$.

$$h_t = \phi(W_{hu} u_t + W_{hh} h_{t-1} + b_{hu}) \tag{4}$$

$W_{hh} \in \mathbb{R}^{d_h \times d_h}$, $W_{hu} \in \mathbb{R}^{d_h \times d_u}$, $b_{hu} \in \mathbb{R}^{d_h}$, represent the RNN's weights to be optimized and $d_h$, $d_u$ are the dimensions of $h_t$ and $u_t$ vectors respectively. Note that the weights are shared across the network. The use of RNN allows the model to learn long-range dependencies where the network is unfolded as many times as the length of the sequence (i.e. length of original and mutated sequences) it is modeling. Although RNNs are capable of handling and representing variable-length sequences, in practice, the learning process faces challenges due to the vanishing/exploding gradient problem [5–7]. In this work, we used gated recurrent unit (GRU) [3, 4] to overcome the latter challenges by updating the computation mechanism of the hidden state vector $h_t$ through the specified equations below.

$$\begin{aligned}
z_t &= \sigma(W_{hu}^z u_t + W_{hh}^z h_{t-1} + b_{hu}^z) && \text{(update gate)} \\
r_t &= \sigma(W_{hu}^r u_t + W_{hh}^r h_{t-1} + b_{hu}^r) && \text{(reset gate)} \\
\tilde{h}_t &= \phi(W_{hu}^{\tilde{h}} u_t + r_t \odot W_{hh}^{\tilde{h}} h_{t-1} + b_{hu}^{\tilde{h}}) && \text{(new state/memory cell)} \\
h_t &= (1 - z_t) \odot \tilde{h}_t + z_t \odot h_{t-1} && \text{(hidden state vector)}
\end{aligned}$$

The GRU model computes a reset gate $r_t$ that is used to modulate the effect of the previous hidden state vector $h_{t-1}$ when computing the new memory vector $\tilde{h}_t$. The update gate $z_t$ determines the importance/contribution of the newly generated memory vector $\tilde{h}_t$ compared to the previous hidden state vector $h_{t-1}$ when computing the current hidden vector $h_t$. The weights $W_{hu}^z$, $W_{hu}^r$, $W_{hu}^{\tilde{h}}$ each $\in \mathbb{R}^{d_h \times d_u}$ and $W_{hh}^z$, $W_{hh}^r$, $W_{hh}^{\tilde{h}}$ each $\in \mathbb{R}^{d_h \times d_h}$. The biases $b_{hu}^z$, $b_{hu}^r$, $b_{hu}^{\tilde{h}}$ each $\in \mathbb{R}^{d_h}$ where $d_h$ and $d_u$ are the dimensions of $h_t$ and $u_t$ vectors respectively. The operator $\sigma$ represents the *sigmoid* function, $\phi$ the *tanh* or *ReLU* function, and $\odot$ the element-wise product (i.e. Hadamard product).

In our work we used a bidirectional GRU that computes two hidden state vectors $\overrightarrow{h_t}$ and $\overleftarrow{h_t}$ for each element $u_t$ in sequence $\underline{U}$ corresponding to left-to-right and right-to-left GRU encoding of the sequence at position $t$. Hence, the newly computed sequence $\underline{G} = [\overleftrightarrow{g_1}, \overleftrightarrow{g_2}, \cdots, \overleftrightarrow{g_T}]$ is based on the *concatenation* of the computed hidden vector representations at each position $t$ (see Eq. 5)

$$\overleftrightarrow{g_t} = [\overrightarrow{h_t} \bigoplus \overleftarrow{h_t}] \tag{5}$$

where $\overleftrightarrow{g_t} \in \mathbb{R}^{d_g}$ and $d_g = 2 \times d_h$.

##### 1.2.3 Attention layer block

The attention layer served as a *pooling* or *compression* layer where the sequence of learned representations $\underline{G}$ are mapped to one fixed-vector representation. We adapted the idea of a *global* attention model [8,9] in which a global context/query vector $q_{global}$ (i.e. trainable parameters) was used along with the output $\underline{G} = [\overleftrightarrow{g_1}, \overleftrightarrow{g_2}, \cdots, \overleftrightarrow{g_T}]$ from the sequence encoder block to generate a pooled representation vector $z_{global}$ (Eq. 6). The objective is to compute *attention* weights $\alpha_t$ for every $\overleftrightarrow{g_t}$ vector where $\alpha_t$ is the normalized weight computed using Eq. 7.

$$z_{global} = \sum_{t=1}^{T} \alpha_t \overleftrightarrow{g_t} \tag{6}$$

$$\alpha_t = \frac{\exp\left(score(q_{global}, \overleftrightarrow{g_t})\right)}{\sum_{k=1}^{T} \exp\left(score(q_{global}, \overleftrightarrow{g_k})\right)} \tag{7}$$

Similarly, we used another attention pooling layer comprising of a trainable context vector $q_{RTT}$ (i.e. trainable parameters) focused on the RTT region of the sequence where mutations are encoded. As a result, the attention weights were computed only based on the elements belonging to the RTT region and later were used to generate a weighted representation $z_{RTT}$ of these elements

$$z_{RTT} = \sum_{t=1}^{T} (\alpha_t \overleftrightarrow{g_t}) \cdot \mathbb{1}[p_t^{RTT} = 1] \tag{8}$$

where $\alpha_t$ is the normalized weight computed using Eq. 9, and $\mathbb{1}[p_t^{RTT} = 1]$ is an indicator variable equal to 1 when the element at position $t$ is part of the RTT region and 0 otherwise.

$$\alpha_t = \frac{\exp\left(score(q_{RTT}, \overleftrightarrow{g_t}, \mathbb{1}[p_t^{RTT} = 1])\right)}{\sum_{k=1}^{T} \exp\left(score(q_{RTT}, \overleftrightarrow{g_k}, \mathbb{1}[p_k^{RTT} = 1])\right)}. \tag{9}$$

For the attention *scoring* function, we defined *score* using the scaled dot-product as in [10] (see Equations 10 and 11 corresponding to the global vector $q_{global}$ and RTT vector $q_{RTT}$ respectively). In both equations, the score is computed by performing a *dot-product* between the query vector $q$ and $\overleftrightarrow{g_t}$ scaled by $d_g$ which is the dimension of both vectors.

$$score(q_{global}, \overleftrightarrow{g_t}) = \frac{q_{global}^{\top} \cdot \overleftrightarrow{g_t}}{\sqrt{d_g}} \tag{10}$$

$$score(q_{RTT}, \overleftrightarrow{g_t}, \mathbb{1}[p_t^{RTT} = 1]) = \frac{q_{RTT}^{\top} \cdot \overleftrightarrow{g_t}}{\sqrt{d_g}} \cdot \mathbb{1}[p_t^{RTT} = 1] \tag{11}$$

After the attention layer operations, the sequence encoder block generates a pair of vectors $z_{global}$ and $z_{RTT}$ for each of the original and mutated sequence separately (i.e. totaling four vectors) that are stacked to create $z_{seq}$ (Eq. 12)

$$z_{seq} = [z_{global}^{original} \bigoplus z_{RTT}^{original} \bigoplus z_{global}^{mutated} \bigoplus z_{RTT}^{mutated}] \tag{12}$$

#### 1.3 Sequence-level features encoder network

In addition to using the original and mutated sequences, we further derived *sequence-level features* such as the melting temperature or minimum free energy for the different parts of the sequences. We derived 18 features that were passed as an input vector to a *sequence-level encoder* comprising a stack of multilayer feed-forward networks, residual connections and layer-normalization operations to map the input features to a dense vector representation. Formally, an input vector $f_{seq} \in \mathbb{R}^{d_f}$ representing *sequence-level features* is mapped using an affine transformation matrix (Eq. 13)

$$f'_{seq} = W_f f_{seq} + b_f \tag{13}$$

where $W_f \in \mathbb{R}^{d' \times d_f}$, $b_f \in \mathbb{R}^{d'}$, $d_f$, and $d'$ representing the dimensions of $f_{seq}$ and $f'_{seq}$ vectors respectively. The embedded vector $f'_{seq}$ is later passed to a feed-forward network consisting of two affine transformation matrices and non-linear activation function to further compute/embed a new vector representation. The first transformation (Eq. 14) uses $W_{MLP1} \in \mathbb{R}^{\xi d' \times d'}$ and $b_{MLP1} \in \mathbb{R}^{\xi d'}$ to transform input $f'_{seq}$ to new vector $\in \mathbb{R}^{\xi d'}$ where

$\xi \in \mathbb{N}$ is multiplicative factor. A non-linear function such as $ReLU(z) = max(0, z)$ is applied followed by another affine transformation using $W_{MLP2} \in \mathbb{R}^{d' \times \xi d'}$ and $b_{MLP2} \in \mathbb{R}^{d'}$ to obtain vector $\tilde{f}_{seq} \in \mathbb{R}^{d'}$. A layer normalization (Eq. 15) is applied to obtain $\tilde{f}'_{seq} \in \mathbb{R}^{d'}$.

$$\tilde{f}_{seq} = W_{MLP2} ReLU(W_{MLP1} f'_{seq} + b_{MLP1}) + b_{MLP2} \quad (14)$$

$$\tilde{f}'_{seq} = LayerNorm(\tilde{f}_{seq}) \quad (15)$$

Layer normalization [11] was used with the goal to ameliorate the "covariate-shift" problem by re-standardizing the computed vector representations (i.e. using the mean and variance across the features/embedding dimension $d'$). Given a computed vector $\tilde{f}_{seq}$, $LayerNorm$ function will standardize the input vector using the mean $\mu$ and variance $\sigma^2$ along the features dimension $d'$ and apply a scaling $\gamma$ and shifting step $\beta$ (Eq. 18). $\gamma$ and $\beta$ are learnable parameters and $\epsilon$ is small number added for numerical stability.

$$\mu = \frac{1}{d'} \sum_{j=1}^{d'} \tilde{f}_{seq(j)} \quad (16)$$

$$\sigma^2 = \frac{1}{d'} \sum_{j=1}^{d'} (\tilde{f}_{seq(j)} - \mu)^2 \quad (17)$$

$$LayerNorm(\tilde{f}_{seq}) = \gamma \times \frac{\tilde{f}_{seq} - \mu}{\sqrt{\sigma^2 + \epsilon}} + \beta \quad (18)$$

Furthermore, we used residual connections / skip-connections [12] in order to improve the gradient flow in layers during training. This is done by summing both the newly computed output of the current layer with the output from the previous layer. In our setting, a residual connection is applied by summing $f'_{seq}$ vector (first embedded vector) with the $\tilde{f}_{seq}$ vector and then passed to a layer normalization layer.

#### 1.4 Decoder block

The last layer in the model takes as input the computed representation vectors from the two **sequence encoder blocks** (original and mutated sequence) $z_{seq}$ (Eq. 12), and the **sequence-level features encoder block** $\tilde{f}'_{seq}$. These vectors are concatenated to become one input vector $z_{final}$

$$z_{final} = [z_{global}^{original} \bigoplus z_{RTT}^{original} \bigoplus z_{global}^{mutated} \bigoplus z_{RTT}^{mutated} \bigoplus \tilde{f}'_{seq}] \quad (19)$$

$z_{final}$ is passed through a stack of multilayer feed-forward networks, residual connections and layer-normalization operations to compute a probability distribution on the outcomes (i.e. **edited**, **unedited**, **unintended edits**). An affine transformation matrix is first applied (Eq. 20)

$$z'_{final} = W_z z_{final} + b_z \quad (20)$$

where $W_z \in \mathbb{R}^{d' \times d_z}$, $b_z \in \mathbb{R}^{d'}$, $d_z$, and $d'$ representing the dimensions of $z_{final}$ and $z'_{final}$ vectors respectively. The embedded vector $z'_{final}$ is later passed to a feed-forward network consisting of two affine transformation matrices and non-linear activation function to further compute/embed a new vector representation. The first transformation (Eq. 21) uses $W^z_{MLP1} \in \mathbb{R}^{\xi d' \times d'}$ and $b^z_{MLP1} \in \mathbb{R}^{\xi d'}$ to transform input $z'_{final}$ to new vector $\in \mathbb{R}^{\xi d'}$ where $\xi \in \mathbb{N}$ is multiplicative factor. A non-linear function such as $ReLU(z) = max(0, z)$ is applied followed by another affine transformation using $W^z_{MLP2} \in \mathbb{R}^{d' \times \xi d'}$ and $b^z_{MLP2} \in \mathbb{R}^{d'}$ to obtain vector $\tilde{z}_{final} \in \mathbb{R}^{d'}$. A layer normalization (Eq. 22) is applied to obtain $\tilde{z}'_{final} \in \mathbb{R}^{d'}$.

$$\tilde{z}_{final} = W^z_{MLP2} ReLU(W^z_{MLP1} z'_{final} + b^z_{MLP1}) + b^z_{MLP2} \quad (21)$$

$$\tilde{z}'_{final} = LayerNorm(\tilde{z}_{final}) \quad (22)$$

A last affine transformation is applied to $\tilde{z}'_{final}$ followed by *softmax* operation to compute a probability distribution on the outcomes (i.e. three outcomes in our current setting). That is, the probability distribution $\hat{y}$ is computed using Eq. 23

$$\hat{y} = \sigma(W_o \tilde{z}'_{final} + b_o) \tag{23}$$

where $\hat{y} \in \mathbb{R}^{|Y|}$, $W_o \in \mathbb{R}^{|Y| \times d'}$, $b_o \in \mathbb{R}^{|Y|}$, $Y$ is the set of admissible outcomes, $|Y|$ is the number of outcomes (i.e. proportions of three outcomes in our case), $d'$ is the dimension of $\tilde{z}'_{final}$ and $\sigma$ is the *softmax* function.

#### 1.5 Objective Function

We experimented with cross-entropy and Kullback–Leibler divergence ($D_{KL}$) as a loss function for an $i$-th design variant sample (i.e. pair of original and mutated sequences). $D_{KL}$ represents the relative entropy of the model's estimated distribution over all target outcomes $\hat{y}$ with respect to the true distribution $y$ (i.e. observed proportions of edited, unedited and unintended edits) (Eq. 24).

$$D^i_{KL}(y^i || \hat{y}^i) = \sum_{j=1}^{|Y|} y^i_j log(\frac{y^i_j}{\hat{y}^i_j}) \tag{24}$$

Lastly, the objective function for the whole training set is defined by the average loss across all the design variant sequences plus a weight regularization term $\lambda$ (i.e. $l_2$-norm regularization) applied to the model parameters represented by $\boldsymbol{\theta}$

$$L(\boldsymbol{\theta}) = \frac{1}{N} \sum_{i=1}^{N} D^i_{KL}(y^i || \hat{y}^i) + \frac{\lambda}{2} ||\boldsymbol{\theta}||^2_2 \tag{25}$$

In practice, the training occurs using mini-batches where computing the loss function and updating the parameters/weight occur after processing each mini-batch of the training set.

#### 1.6 Multi-task learning with multiple datasets

Conventional approaches have often proposed training separate models for each dataset (i.e cell line). In this work, we propose a multi-task learning model that leverage common patterns and relationships present across various datasets (i.e. applied to different cell lines) instead of training separate models for each dataset. The goal is to train one model that exploits the common patterns across datasets while reducing the computational run time and model size without sacrificing the prediction performance.

Given a total number of $S$ datasets where each dataset $S_i$ (denoted by screening library), we developed a multi-task learning model that uses shared encoding layers to extract a common representation across all the libraries as well as individual branches that train the model specifically for each library. We experimented with multiple options (Table 1) where we change the set of shared and separate blocks when training across all datasets. For our final models, we opted for option 3 that uses a shared embedding and encoding layers and creates a separate set of blocks for the attention, sequence level feature embedding and decoder layers for each dataset. To counterbalance any bias towards larger datasets, we implemented a data loader that uniformly samples the same number of data samples in each mini-batch throughout the training phase.

| | Embedding layer | Encoder layer (bidirectional RNN + other embedding blocks) | Attention layer | Sequence-level feature embedding layer | Decoder layer |
|---|---|---|---|---|---|
| Option 1 | Shared | Shared | Shared | Shared | Separate |
| Option 2 | Shared | Shared | Separate | Shared | Separate |
| Option 3 | Shared | Shared | Separate | Separate | Separate |

Table 1: Multi-task learning model design options

#### 1.7 Pretraining & Fine-tuning models

We also experimented with pretraining models on large datasets and then fine-tuning them on our 'Library-Diverse' (our current dataset from HEK293T and K562 cell lines). This translates to first training a **base model** denoted by **Model A (PRIDICT1.1)** on a large dataset such as 'Library 1' (92,423 pegRNAs, from Mathis et al. 2023 [1]), then fine-tuning this model in multi-task learning setup (jointly learning on both datasets from 'Library-Diverse'). In our setting, fine-tuning is defined by freezing a chosen set of weights of the pretrained

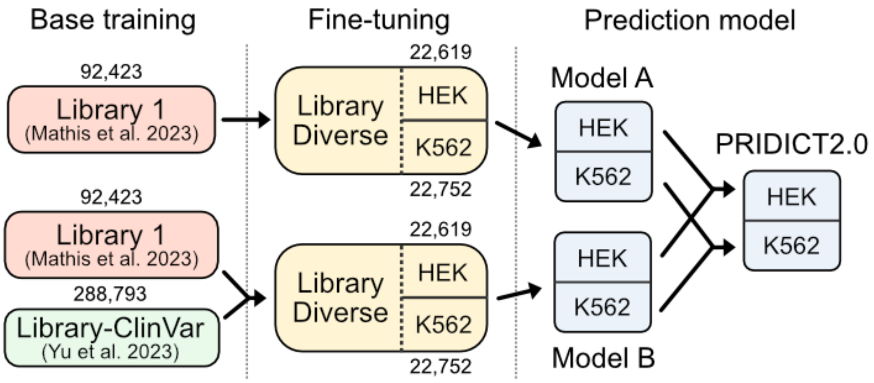


Figure 1: Overview of the trained models

model (i.e. **Model A (PRIDICT1.1)**) and then retraining the rest of the model weights. We decided to freeze the weights corresponding to the shared layers described in option 3 (Table 1) and retrain the ones that are in the separate layers (i.e. dataset specific layers). In other words, the embedding and encoder layers were frozen (i.e. using the weights from pretrained model) and the rest are retrained using multi-task learning. We repeated this process by creating another **base model** denoted by **Model B (PRIDICT1.2)** that is trained using both 'Library 1' and 'Library-ClinVar' (288,793 pegRNAs, from Yu et al. 2023 [13]). This base model was trained in a multi-task learning setup on both datasets to create the pretrained base model. Then we proceeded to fine-tune it on our 'Library-Diverse'. Figure 1 shows an overview of the trained models.

### Supplementary Methods 1b: Experiments for machine learning

#### 2.1 Training & Evaluation Workflow

Our 'Library-Diverse' dataset included 22,619 (HEK293T) and 22,752 (K562) pegRNAs that were used to train and evaluate machine learning models. For all models, we followed a grouped 5-fold cross-validation on Library-Diverse where pegRNAs for the same locus were kept in the same train- or test set. Each fold had 80% of the Library-Diverse pegRNAs for training and 20% for testing. A validation set for each fold was created by taking a 10% grouped random split from the fold's training sequences that was used to optimize model hyperparameters.

We trained three models with different setups. The first model denoted by **PRIDICT-LibDiv** was trained on 'Library Diverse' using multi-task learning setup (option 3 in Table 1). For the second model, we started with pretraining a base model on 'Library 1' and then fine-tuning it on our 'Library Diverse' to create **Model A (PRIDICT1.1)** again following multi-task learning (option 3 in Table 1). For the third model, we first pretrained a base model on 'Library 1' and 'Library-ClinVar' datasets, then fine-tuned on 'Library-Diverse' to create **Model B (PRIDICT1.2)**. All models are based on **PRIDICT** architecture and when multiple datasets ($\geq 2$) were involved in training, multi-task learning setup was used. Finally, **Model A (PRIDICT1.1)** and **Model B (PRIDICT1.2)** were combined to form an ensemble model **PRIDICT2.0** by combining prediction values in a 1:1 ratio.

We evaluated models' performance using Pearson and Spearman correlation. During models' training, the epoch in which the model achieved the best harmonic mean between both scores on the validation set was recorded, and model state as it was trained up to that epoch was saved. This best model, as determined by the validation set, was then tested on the test split. The evaluation of the trained models was based on their average performance on the test sets across the five folds.

#### 2.2 Hyperparameters Optimization

We used a uniform random search strategy [9, 14] that randomly chose a set of hyperparameters configurations (i.e. embedding dimension, number of hidden layers, dropout probability, etc.) from the set of all possible configurations and trained corresponding models on a random fold. Then the best configuration for each model (i.e. the one achieving best performance on the validation set) was used for the final training and testing of each model on all 5 folds.

The range of possible hyperparameters configuration (i.e. choice of values for hyperparameters) for PRIDICT models is reported in Table 1.

**List 1** PRIDICT hyperparameters options

- `Embedding Block operations`
  - `Nucleotide embedding layer`
    - `embedding dimension` $d'$ ........ $\{16, 32, 64, 128\}$
  - `Annotation embedding layer`
    - `embedding dimension` $d_a$ ........ $\{4, 8, 16\}$
  - `concatenate option` ........ $\{stacking, adding\}$
- `Encoder Block operations`
  - `GRU`
    - `number of hidden layers` ........ $\{1, 2\}$
    - `hidden dimension` ........ $\{16, 32, 64, 128\}$
    - `embedding dimension` ........ $\{16, 32, 64, 128\}$
    - `dropout` ........ $\{0.15, 0.25, 0.35\}$
  - `Sequence-level features encoder`
    - `Feed-Forward (MLP) block`
      - `embedding dimension` ........ $\{16, 32, 64, 128\}$
      - `MLP embedding factor (multiplier)` $\xi$ ........ $\{1, 2, 3\}$
      - `Non-linear function` ........ $\{ELU, ReLU\}$
      - `dropout` ........ $\{0.15, 0.25, 0.35\}$
    - `number of repeats for MLP Block` ........ $\{1, 2\}$
- `Decoder Block operations`
  - `Feed-Forward (MLP) block`
    - `embedding dimension` ........ $\{16, 32, 64, 128\}$
    - `MLP embedding factor (multiplier)` $\xi$ ........ $\{1, 2, 3\}$
    - `Non-linear function` ........ $\{ELU, ReLU\}$
    - `dropout` ........ $\{0.15, 0.25, 0.35\}$
  - `number of repeats for MLP Block` ........ $\{1, 2\}$
- $l_2$`-norm regularization` $\lambda$ ........ $\{10^{-5}, 10^{-4}, 10^{-3}, 10^{-2}, 10^{-1}\}$
- `Batch size during training` ........ $\{1000, 1500, 2000\}$
- `Optimization algorithm` ........ $\{Adam\}$
